# Functional activity of antisera against recombinant Zika virus envelope protein subunits expressed in *Escherichia coli*

**DOI:** 10.1101/698266

**Authors:** Hong-Yun Tham, Man Kwan Ooi, Vinod RMT Balasubramaniam, Sharifah Syed Hassan, Hong-Wai Tham

## Abstract

The global Zika virus (ZIKV) outbreak across continents has been drawing research attentions to researchers and healthcare professionals. It highlights the urgent development of ZIKV vaccines that offer rapid, precise and specific protection to those living in the high-risk regions - the tropical and subtropical regions. As a public health priority, there is a progressive development in the discovery of vaccine candidates and design in recent years. Many efforts have been placed in the *in vitro* development of ZIKV subunits as the vaccine candidate in various protein expression systems, including bacteria, yeast, plant cells, insect cells and mammalian cells. However, due to the lack of knowledge on humoral and cellular immune responses against virus vaccines, a commercialised vaccine against Dengue virus (DENV) has been suspended due to a health scare in Philippines. Moreover, the closely-related DENV and ZIKV has indicated serological cross-reactivity between both viruses. This has led to greater attentions to precautions needed during the design of ZIKV and DENV vaccines. In this study, we pre-selected, synthesised and expressed the domain III of ZIKV envelope protein (namely rEDIII) based on a previously-established report (GenBank: AMC13911.1). The characteristics of purified ZIKV rEDIII was tested using SDS-PAGE, Western blotting and LC-MS/MS. Since the ZIKV rEDIII has been well reported as a potential protein candidate in ZIKV vaccine development, we assessed the possible outcome of preexisting immunity against the rEDIII proteins by conducting dot-blotting assays using mice antisera pre-immunised with ZIKV particles (ZIKV strain: MRS_OPY_Martinique_PaRi_2015, GenBank: KU647676). Surprisingly, the antisera was able to recognise the rEDIII of a different ZIKV strain (GenBank: AMC13911.1). Despite its great antigenicity in eliciting humoral and cellular immunity against ZIKV infection, our finding calls for greater attention to evaluate the details of ZIKV rEDIII as a stand-alone vaccine candidate.

## Introduction

Zika virus (ZIKV), a member of the *Flaviviridae* family, is transmitted between humans by its main mosquito vectors, *Aedes aegypti* or *Aedes albopictus* [1, 2]. ZIKV carries a single-stranded, positive-strand RNA genome of about 11 kb in length [3]. Despite its first isolation from *Rhesus macaque* in 1947, limited reports on human infection is available. This is largely due to its self-limited illnesses including low-grade fever, headache, myalgia and arthralgia [4–6]. There are two lineages (African and Asian) and three genotypes (East African, West African, and Asian) of ZIKV circulated in tropical and subtropical regions [7]. As of 2018, the diagnosis assays for ZIKV comprised of 5 serological assays and 14 molecular assays with Food and Drug Administration Emergency Use Authorisation (FDA EUA), which have been well reviewed by in 2018 by Theel and Hata [8]. On the other hand, in terms of vaccine candidate discovery and development, a recent article by Alan Barett reported that over 45 vaccine candidates have been discovered, with at least 9 are currently in clinical evaluation [9]. Nevertheless, it is not plausible to develop an efficacious ZIKV vaccine in near future due to the serological cross reactivity of antibodies between Dengue virus (DENV) and ZIKV [10–14]. In addition, similarities in transmission process, disease manifestations and transmitting vectors between Zika fever and Dengue fever are often confused [15, 16], which have further halted the development of ZIKV-specific vaccine candidate.

Since its declaration as Public Health Emergency of International Concern by World Health Organisation (WHO) in February 2016, Zika virus has been associated with microcephaly and neurological complications such as Guillain-Barré syndrome [17, 18]. Since then, international attention has been brought towards the rapid chain of disease outbreak, which was spread throughout South, Central and part of North America, followed by Asia Pacific [19, 20]. Vector transmission of Zika fever occurs mainly in tropical regions. However, cases in returning travellers have frequently been reported in locations including Europe, US, Australia, New Zealand, Japan, UK, and China [21–25].

Upon infection, ZIKV was reported to persist in body fluids, such as urine or saliva, for longer than that of in the blood [26–28]. This becomes an important consideration in the development of rapid and effective tools for ZIKV detection. Prior to 2018, several research groups reported various ZIKV detection strategies, including a newly developed strategy - liposome-based immunoassay reported by Shukla *et. al*. [29], who reported the low sensitivity of 5 commercially available immunoassays to detect ZIKV infection [30]. Soon after, Powley *et. al*. reviewed the current methods of ZIKV detection and their limitations [31]. The authors highlighted a few restrictions including the need of expensive machineries, trained personnels, intensive laborious processes, viral RNA stability, lack of specific anti-ZIKV antibodies, and possibilities of false-positive results with current diagnostic techniques. In addition, Pawley *et. al*. also emphasised the importance of anti-ZIKV monoclonal antibodies in the development of novel point-of-care paper-based detection method [31]. This has again emphasised the importance of the domain III of ZIKV envelope protein, which carries a strong antigenicity and greatest power of discrimination from other members of Flavivirus [13, 32], as an ideal protein candidate in ongoing and future development of point-of-care testing for active infection for ZIKV.

Recombinant domain III of ZIKV envelope protein (rEDIII) has been previously expressed and purified using different protein expression systems, including yeast [33], insect cells [34, 35], plant cells [36, 37] and bacterial cells [34, 35, 38]. Sylvia *et. al*. proved the integrity of rEDIII through SDS-PAGE, Western blot and immunoblotting [39], while Yang *et. al*. described the generation and immunogenicity of the ZIKV rEDIII as a protein subunit vaccine candidate, which was also demonstrated to elicit anti-rEDIII monoclonal antibody in pre-clinical studies [35]. In accordance with this, this study was designed not only to construct a protein-expression plasmid for recombinant ZIKV envelope protein (domain III, rEDIII) production, but also to assess the possibilities of the ZIKV rEDIII to cause antibody dependent enhancement (ADE) or serum sickness (SS) in the recipients, especially those who had exposed to ZIKV infection prior to receiving vaccine which contains rEDIII as the vaccine candidate.

## Materials and methods

### Zika virus EDIII gene

Complete coding sequence of the domain III of Zika virus (strain: PRVABC59) envelope protein (EDIII) was retrieved from National Centre for Biotechnology Information (NCBI) (GenBank accession number: AMC13911.1) [35, 36]. Gene block and primers were synthesised (Integrated DNA Technologies, IDT^®□^) and stored in −20 °C until used.

### Gene cloning and protein expression

ZIKV EDIII gene was synthesised and cloned in pUCIDT plasmid vector, namely pUCIDT-ZVEDIII. The plasmid was transformed into *Escherichia coli* DH5α strain for long-term storage at −80 °C. After plasmid purification, ZIKV EDIII coding sequence was amplified using primers (forward: TCTGCAGCTGGTACCGCGTTCACATTCACCAAGATCCCGGCTG; reverse: TCAAGCTTCGAATTCTGCTTTTCCAATGGTGCTGCCACTCCTG) with the following PCR conditions: 1 cycle of 94 °C (2 minutes); 35 cycles of 94 °C (45 seconds), 55 °C (45 seconds), 72 °C (1 minute); 1 cycle of 72 °C (10 minutes); on hold at 4 °C until use. PCR product was cloned in-frame into pRSET-B protein expression vector (Invitrogen, CA, USA) using In-Fusion^®^□ HD Cloning Plus (Takara Bio, USA). Recombinant plasmid was transformed into competent *E. coli* BL21 (DE3) strain for protein expression analysis.

For protein expression, an overnight culture of transformed BL21 (DE3) *E. coli* was diluted to 1:100 with Luria Bertani broth supplemented with ampicillin at final concentration of 75 μg/mL. Bacteria culture was incubated (37 °C, 180 rpm) until OD_600_ of 0.50 was reached. Protein expression was induced by the addition of IPTG to the final concentration of 1 mM and incubation was further conducted for 3 hours (37 °C, 180 rpm). Then, the cells were harvested by centrifugation (3000 *g*, 4 °C, 2 minutes). Cell pellet was resuspended in SDS reducing buffer, aliquoted into 50 μL, heated at 99 °C for 10 minutes before loading into a 12% SDS acrylamide gel.

### SDS-PAGE and Western blot

SDS-PAGE was conducted in vertical direction at 100 V in a 1x Tris-glycine running buffer (25 mM Tris, 192 mM Glycine, 0.1% SDS, pH 8.3) [40]. After that, protein bands were stained with R-250 Coomassie Brilliant Blue stain. Another duplicated gel was subjected to Western blotting. Protein bands were transferred onto PVDF membrane [41], followed by blocking (5% BSA, 1 hour, 25 °C), primary antibody (anti-Xpress monoclonal antibody, 1:5000 dilution, 1 hour, 25 °C), and secondary antibody (anti-mouse IgG, 1:5000 dilution, 1 hour, 25 °C). Protein bands were visualised by addition of substrate (BCIP/NBT).

### rEDIII purification

ZIKV rEDIII was purified with gradual decrease of urea concentration (8 M, 6 M, 4 M, 2 M and 0 M) to progressively remove urea through dialysis. All buffers (except elution buffer) were supplemented with 20 mM imidazole to reduce nonspecific binding of unwanted protein to the HisTrap HP histidine-tagged protein purification columns (GE Healthcare). In brief, after protein expression, cell pellets of transformed BL21 (DE3) *E. coli* was suspended in dissolving buffer supplemented with 8 M urea. Mixture was incubated in HisTrap HP histidine-tagged protein purification columns at room temperature for 30 minutes, followed by washing steps using a series of buffers supplemented with 6 M, 4 M, 2 M and 0 M of urea. Lastly, rEDIII was eluted with elution buffer (supplemented with 0 M urea and 500 mM of imidazole). Eluents were subjected to dialysis using 1x PBS buffer at 4 °C for 2 hours. Purified rEDIII was kept at 4 °C for further analyses.

### LC-MS/MS

The rEDIII protein band was excised from polyacrylamide gel and the sample was prepared for *de novo* protein sequencing using in-gel digestion according to manufacturer’s protocol (Agilent Technologies, Inc., 2015). Briefly, the excised gel slice was destained with 200 mM of ammonium bicarbonate (ABC) in 40% acetonitrile (ACN), followed by reduction and alkylation by DTT and IAA respectively. After that, gel slice dehydrated by 100% CAN (15 min, 37°C). The dehydrated gel slice was incubated with trypsin (16 hours, 37 °C) and the reaction was stopped by addition of formic acid. The tryptic peptides were further extracted from the gel slices using 50% ACN and 100% ACN for 15 min each. The recovered peptides were analysed using Agilent 1200 HPLC-Chip/MS interface, coupled with Agilent 6550 iFunnel Q-TOF LC/MS. The *de novo* sequences was analysed and aligned using PEAKS 8.0 software [42].

### rEDIII protein integrity test

Gold Syrian hamsters were bred and housed at the specific pathogen free (SPF) animal facilities, Monash University Malaysia. Ethics approval for animal housing and experimentation were obtained (Monash Animal Ethics: MARP/2017/060). Hamsters were administered subcutaneously with Zika virus (strain: MRS_OPY_Martinique_PaRi_2015, NCBI: KU647676) with TiterMax adjuvant at 10^7^ pfu. After 35 days, serum sample were collected to determine its binding ability towards ZIKV rEDIII proteins.

The integrity of ZIKV rEDIII was determined through Dot Blot assay. First, purified rEDIII was immobilised on a PVDF membrane at 1 μg per dot. rEDIII were dried at 25 °C before blocking (5% BSA, 1 hour, 25 °C). After washing, mouse serum (1:500) were applied (1 hour, 25 °C), followed by anti-mouse IgG (1:5000, 1 hour, 25 °C) before visualisation using BCIP/NBT as the substrate. Control spots were also conducted concurrently using mock-infected mouse serum.

## Results

### rEDIII expression

The coding sequence of ZIKV rEDIII (GenBank accession number: AMC13911.1) was synthesised and cloned in-frame into pRSET-B protein expression vector for protein expression using *E. coli* BL21 (DE3). After SDS-PAGE, based on the molecular weight, the rEDIII was expressed at its expected size (total of 14 kDa) with the 11 kDa moiety carrying a 6x histidine tag at the N-terminal of the recombinant protein. Hence the total expected protein size of 14 kDa (Fig 1).

**Fig 1.**
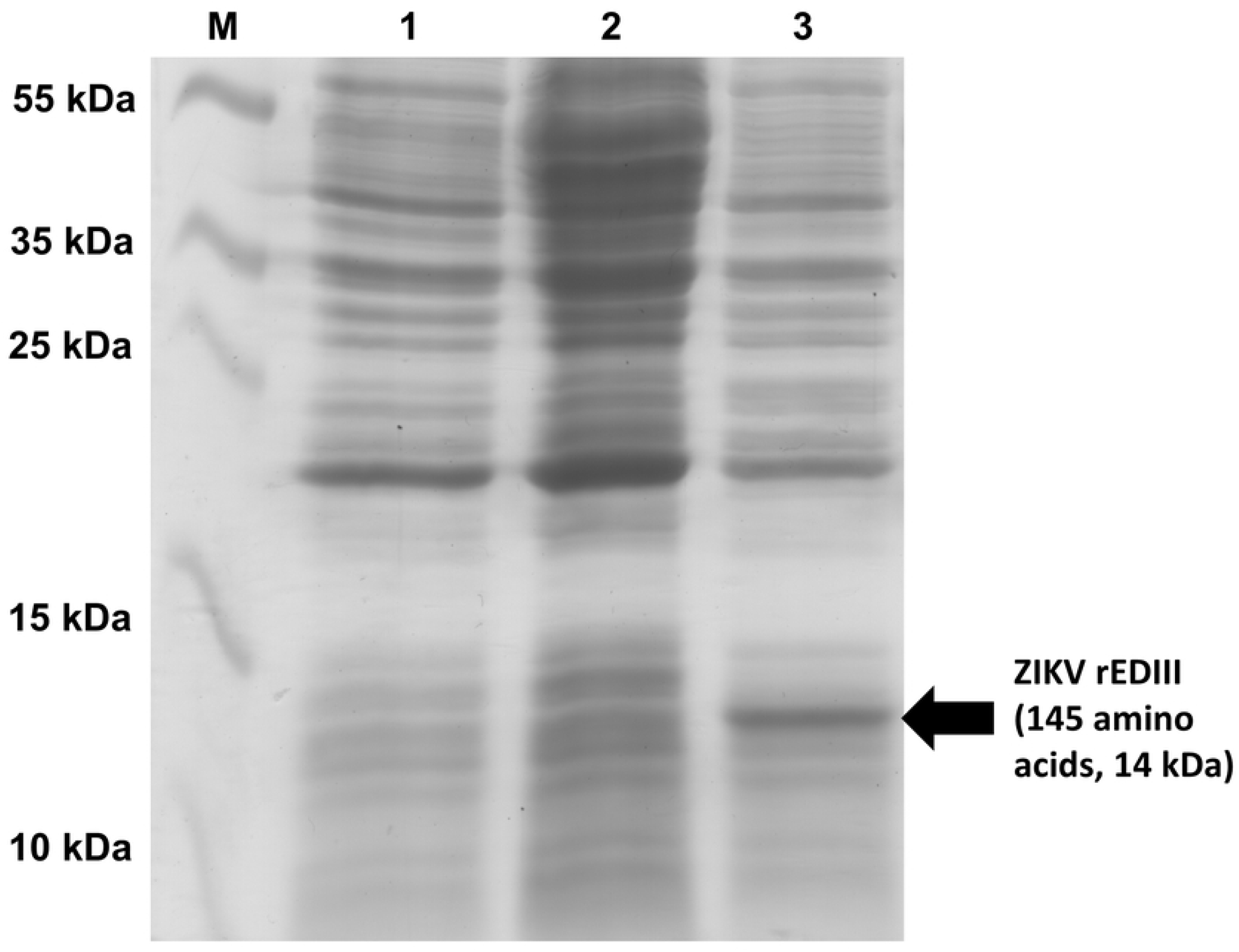
Expression of recombinant domain III of Zika virus (ZIKV) envelope protein (rEDIII) in *Escherichai coli* BL21 (DE3). (M) Protein marker; (1) Negative control: The cellular lysate of untransformed BL21 (DE3) *E. coli*; (2) Supernatant of transformed and IPTG-induced BL21 (DE3) *E. coli* after sonication; (3) Pelleted inclusion body and cellular debri of transformed and IPTG-induced BL21 (DE3) *E. coli* after sonication. The presence of ZIKV rEDIII is indicated by black arrow.

### Western blotting

Western blot was conducted on PVDF membrane using anti-Xpress antibody as the primary antibody. Ther result showed that ZIKV rEDIII was expressed at the expected size (14 kDa) (Fig 2).

**Fig 2.**
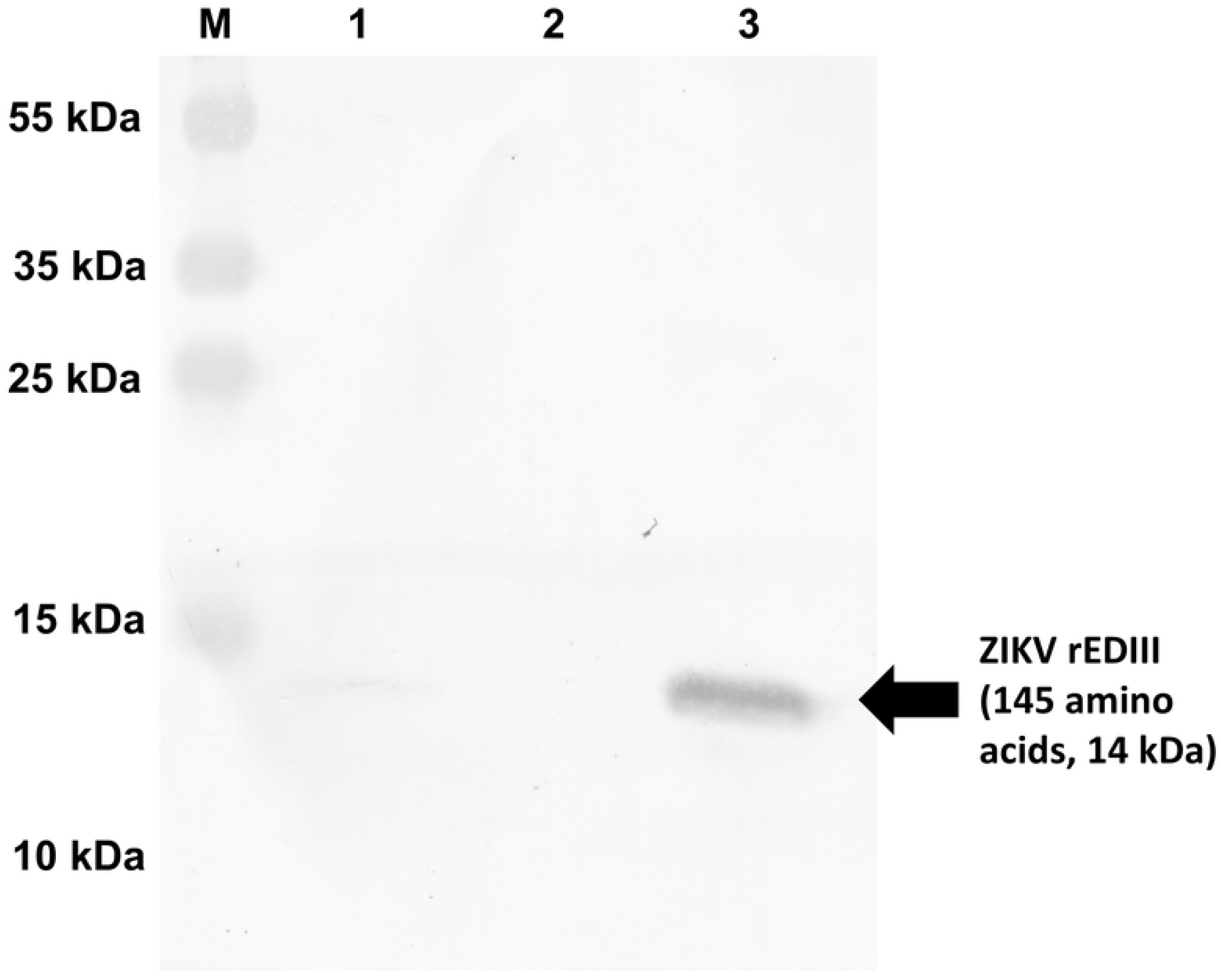
Western blotting analysis shows the presence of the recombinant domain III of Zika virus (ZIKV) envelope protein (rEDIII) at the expected position (14 kDa, indicated by black arrow). (M) Protein marker; (1) Negative control which contains the cellular lysate of untransformed *E. coli* BL21 (DE3); (2) Supernatant of transformed and IPTG-induced *E. coli* BL21 (DE3) after sonication; (3) Pelleted inclusion body and cellular debri of transformed and IPTG-induced *E. coli* BL21 (DE3) after sonication.

### rEDIII purification

ZIKV rEDIII was purified using HisTrap HP histidine-tagged protein purification columns. rEDIII was mainly detected in insoluble inclusion bodies (Fig 3A, lane 2). Lane 2, 3 and 4 was loaded with eluents of washing buffers. The rEDIII was not detected these lanes (Fig 3B, lane 2, 3 and 4). Lastly, purified rEDIII were successfully eluted, which was shown in a single protein band (Fig 3B, lane 5 and 6).

**Fig 3A.**
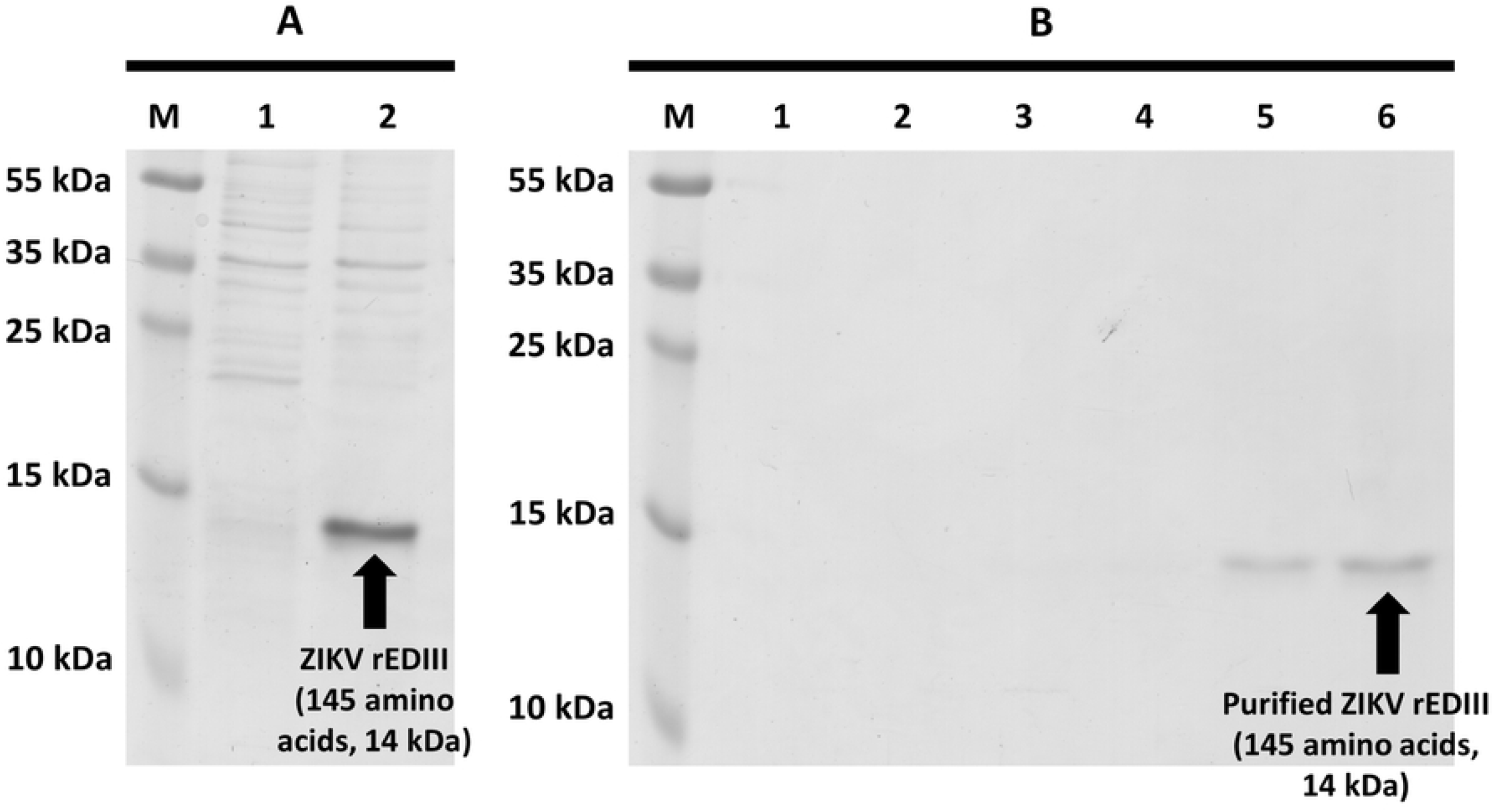
SDS-PAGE analyses of different portions of bacterial cell lysate after IPTG induction. (M) Protein marker; (1) The supernatant of cellular lysate after sonication and centrifugation; (2) The pelleted inclusion body and cell debri after sonication and centrifugation. The distinctive protein band (indicated by black arrow) shows the expected position of ZIKV rEDIII which present in the inclusion body of the *E. coli*. **Fig 3B. The replicated samples of lane 2 in figure 3A were directed to protein purification using Ni-charged resins.** (1) The first flow-through of ZIKV rEDIII inclusion body dissolved in buffers supplemented with 8M urea; (2, 3 & 4) Flow-through of washing buffers supplemented with 6 M, 4 M and 2 M of urea, respectively; (5 & 6) First and second elution of ZIKV rEDIII from the Ni-charged resins using elution buffers supplemented with 500 mM imidazole. ZIKV rEDIII was successfully purified with the expected size of protein indicated by black arrow (14 kDa).

### LC-MS/MS

The sequences of tryptic digested peptides of ZIKV rEDIII were aligned with the protein database through PEAKS DB search and showed alignment with ZIKV polyprotein (A0A0U4ETI0) starting from position 601 to 699 (Fig 4).

**Fig 4.**
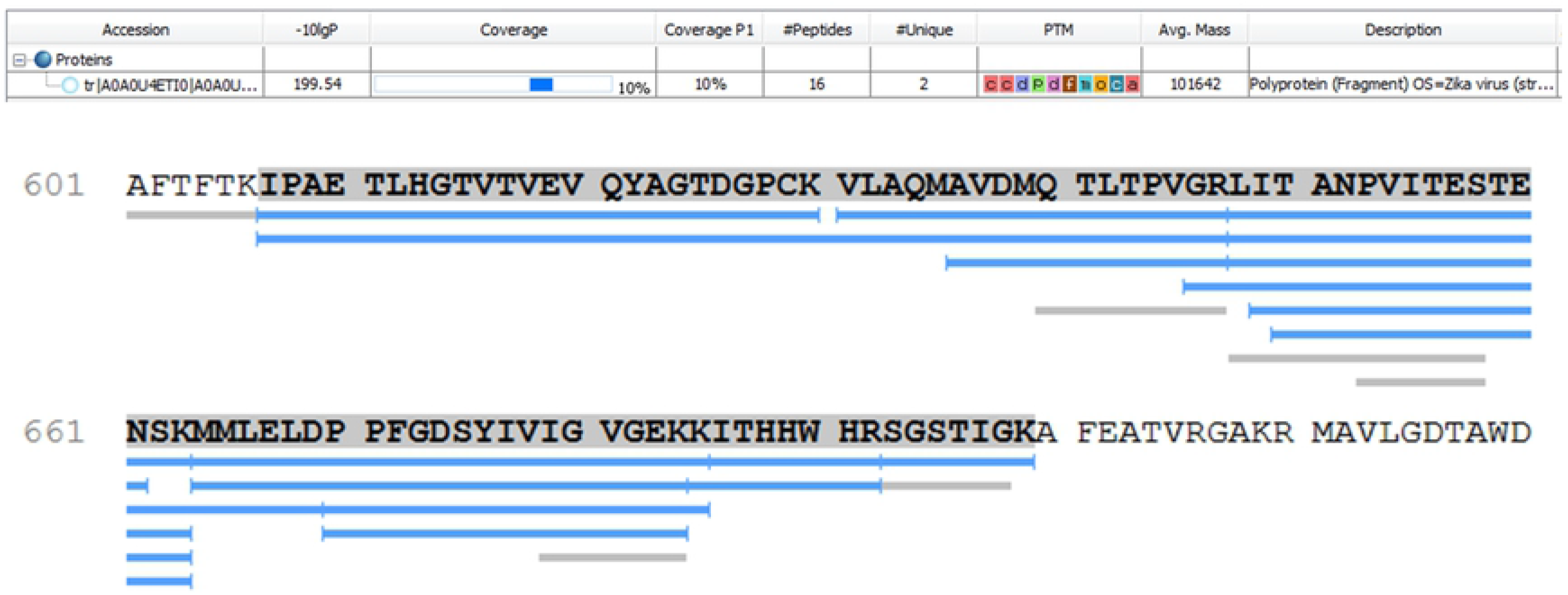
Alignment of tryptic-digested peptides of recombinant ZIKV rEDIII using PEAKS 8.0. The amino acids of subject sequence (domain III of ZIKV) are bold and highlighted in grey. All query sequences are illustrated in blue.

### rEDIII protein integrity test

Dot blot assay was conducted to test the integrity of purified rEDIII. Antisera derived from mock-infected and ZIKV -infected mice and were used. The results showed that antisera was able to recognise the purified rEDIII (Fig 5), which also explains the chances of administered ZIKV rEDIII being recognised by antibodies produced by recipients who were previously infected by ZIKV. On the other hand, no binding to ZIKV rEDIII was observed when the antiserum of mock-infected mice was used in the dot-blot assay.

**Fig 5.**
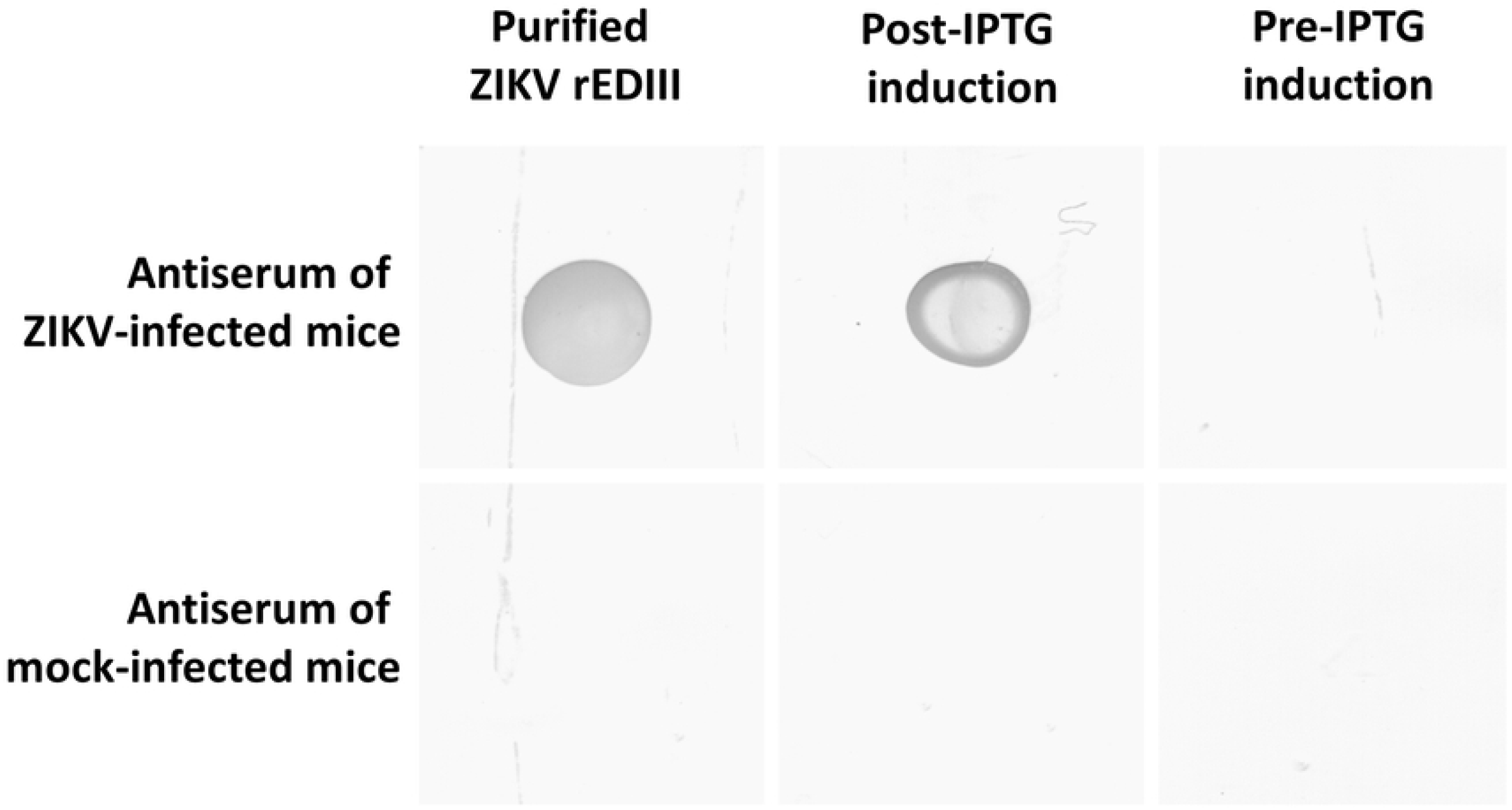
Dot-blot analyses of recombinant domain III of Zika virus (ZIKV) envelope protein (rEDIII). All 6 samples were loaded with either 1 μg of purified ZIKV rEDIII, or bacterial cell lysate of post- or pre-IPTG induction. All samples were air dried prior to incubating with antiserum derived from either ZIKV-infected or mock-infected mice.

## Discussion

Flavivirus is responsible for a number of economically-important diseases in human, including dengue fever, zika fever, yellow fever, West Nile fever and Japanese encephalitis fever. The discovery and development of new vaccine candidates have been in need, and the domain III of flavivirus envelope protein (EDIII) has long been recognised as a suitable candidate due to its high antigenicity and ability to stimulate the production of antibodies by the immune system [43–45]

In this study, we employed the coding sequence of ZIKV rEDIII reported by Yang *et al*. (GenBank accession number: AMC13911.1) in their previous studies for immunogenicity assessments in mammals [35, 36]. Yang et al. concluded that ZIKV rEDIII produced in either plants or *E. coli* had successfully induced immune response to confer sufficient protection against ZIKV infection in mice. However, being one of the most neglected diseases in tropical regions, Zika fever manifested in many individuals without medical attentions, which individuals have developed natural-active immunity against the rEDIII. In order to assess the specificity of these antisera against this ZIKV rEDIII, this study developed and obtained mice antisera containing natural-active antibodies for antibody specificity tests against the ZIKV rEDIII.

A number of concerns have been raised for marketed vaccines, including polio vaccine and measles vaccine which cause toxic shock syndrome [46–49]. Although some avoidable cases were reported to be human-caused [50], the major dengue vaccination programme in Philippines led to a theoretical elevated risk of dengue haemorrhagic fever (DHF) in seronegative vaccine recipients [51]. Although more seroepidemiological surveillance data is needed, adversed manifestations of Zika virus infection due to antibody-dependent enhancement (ADE) has been reported [52, 53]. These data highlighted the possibilities of rEDIII to cause ADE in its recipients.

We infected mice with active virus particles to raise antiserum against ZIKV. In another parallel experiment, the rEDIII was expressed in BL21(DE3) *E. coli*, extracted and purified. The protein identity was confirmed with LC-MS/MS and PEAKS DB search, with the native structure of the rEDIII confirmed by dot blot assays. Meanwhile, the dot blot assay also proved the hypothesis that the antibodies produced by natural active immunity in mammals are able to recognise the our ZIKV rEDIII protein.

ADE caused by administration of vaccine is not uncommon. Understanding immune responses to viral infections is crucial in deciphering the molecular mechanisms behind the enhanced illness by pre-existing antibodies found in the serum of vaccine recipients. Usually, ADE is caused by type III hypersensitivity of the immune system against the vaccine candidate. Cases of ADE after vaccine administration were reported for several vaccine candidates including inactivated and purified influenza virus [54], recombinant Hepatitis B virus [55] and Dengue virus [56, 57]. Since the knowledge and understanding of ADE caused by Zika virus infection is sparse, more attentions should be placed in the development of rEDIII into ZIKV vaccine, where the protein subunit may develop ADE in the vaccine recipients. This is especially important when Dengue virus, the virus that is prevalent in causing ADE, and Zika virus are taxonomically close, with evidences showing serological cross reactivity of antibodies against both viruses in mammals [10–14].

Dejnirattisai *et al*. reported that most of the antibodies against DENV epitopes also bound to ZIKV, but unable to neutralise ZIKV and instead promoted ADE [12]. Recently, other researchers have discussed the risk-to-reward ratio of developing ZIKV vaccine, with regards to current controversial data and unknown interplay between members of flavivirus [53]. These important information must be taken into considerations especially during the development of any virus vaccine.

Several improvements can be made to enhance the bioavailability and effect of ZIKV vaccine using rEDIII as the vaccine candidate. This can be done through the optimisation of adjuvant, which has been thoroughly reviewed by Hogenesch *et al*. in 2018 [58]. HogenEsch *et al*. described the pharmacokinetics of aluminium-based adjuvants, characteristics of antigens, and formulations of vaccines with aluminium adjuvants. On the other hand, in light with its potential wide global distributions of ZIKV vaccine across different continents, the thermodynamic stability of ZIKV rEDIII can also be improved by molecular structural modifications or optimisation of subcellular protein expression [59].

In conclusion, although rEDIII can be the ideal protein candidate in the development of ZIKV vaccine, our results, in conjunction with several previous studies, call for a greater attention on the mechanisms of ADE in vaccine recipients. Our findings are also useful for ZIKV rEDIII applications in the field of virus diagnostics, vaccine developments and viral disease therapies.

## Acknowledgements

Authors acknowledge the continuous support given by the Virus–Host Interaction Research Group (Jeffrey Cheah School of Medicine and Health Sciences, Monash University Malaysia), Biopharmaceutical Research Unit (Faculty of Pharmacy, SEGi University), and Research and Innovation Management Centre (RIMC, SEGi University).

